# Identification of Heterogeneous Cortical Thickness Patterns Associated with Prenatal Gestational Diabetes Exposure: A SuStaIn-Based Subtyping Study

**DOI:** 10.64898/2026.05.25.727436

**Authors:** Yumeng Qi, Eustace Hsu, Seonjoo Lee, Shan Luo, Xi Zhu

## Abstract

**Importance:** Prenatal exposure to gestational diabetes mellitus (GDM) has been associated with adverse metabolic, neurodevelopmental, and psychiatric outcomes in offspring. However, whether GDM-exposed youth exhibit heterogeneous neuroanatomical patterns remains unclear.

**Objective:** To identify distinct cortical thickness subtypes among GDM-exposed youth and examine their associations with anthropometric, neurocognitive, psychiatric/behavioral and neuroimaging measures both cross-sectionally and longitudinally.

**Design, Setting, and Participants:** This cohort study used the Adolescent Brain Cognitive Development (ABCD)®data, a multisite longitudinal population study. Subtype and Stage Inference (SuStaIn), an unsupervised machine learning framework, was applied to cross-sectional structural MRI data to identify cortical thickness patterns in 573 GDM-exposed youth and 2854 healthy controls. Posthoc longitudinal analyses included 1,853 observations from a subset of GDM-exposed youth with 1-, 2-, and 4-year follow-up visits to examine subtype differences in developmental trajectories over time.

**Exposure(s):** Prenatal exposure to GDM.

**Main Outcome(s) and Measure(s):** The primary outcomes included identification of cortical thickness subtypes and their inferred regional ordering patterns. Secondary outcomes included subtype-specific anthropometric, neurocognitive, psychiatric/behavioral and neuroimaging measures.

**Results:** The GDM-exposed sample had a mean age of 119.02 ± 7.34 months and was 47.5% female. Two cortical thickness subtypes were identified. Between subtypes, Subtype 1 (63.2%) was characterized by earlier inferred insula involvement and was associated with greater height (*d* = 0.36, *p*_*FDR*_ < 0.001) and weight (*d* = 0.26, *p*_*FDR*_ = 0.007), whereas Subtype 2 exhibited earlier inferred frontal involvement and nominally higher Attention-Deficit/Hyperactivity Disorder (ADHD) prevalence (*d* = 0.08, *p* = 0.036), steeper longitudinal cortical thinning across all six cortical regions of interest (β range: −0.05 to −0.13, all *p*_*FDR*_ < 0.05), and a smaller decline in Obsessive-Compulsive Disorder (OCD) prevalence over time (β = -1.02, *p*_*FDR*_ = 0.049).

**Conclusions and Relevance:** GDM exposure was associated with two distinct offspring cortical thickness subtypes, each showing different inferred regional ordering patterns and clinical associations. One subtype showed an insula-cingulate–predominant pattern associated with anthropometric measures, whereas the other showed a frontal-predominant pattern associated with nominally higher psychiatric measures and faster cortical thinning over time.

## Introduction

Gestational diabetes mellitus (GDM) is a common medical condition during pregnancy that fundamentally alters the intrauterine metabolic environment. Growing evidence suggests these perturbations have lasting consequences for offspring that extend beyond birth. Offspring exposed to maternal GDM are at significantly increased risk for adverse adiposity-related outcomes, including elevated BMI, higher percent body fat (%BF), and obesity^1–4^. Crucially, this risk profile is not limited to physical health, prenatal GDM exposure has been linked to significant neurodevelopmental and psychiatric vulnerabilities, including elevated risks of Attention-Deficit/Hyperactivity Disorder (ADHD), Autism Spectrum Disorder (ASD), schizophrenia, intellectual disabilities and behavioral problems^5–11^.

Neuroimaging offers insights into the neural correlates associated with prenatal metabolic exposures and their relation to downstream outcomes in offspring. Youth exposed to GDM demonstrate widespread alterations in regional brain morphology^11–15^, functional activation^16–18^, and white-matter microstructure^19,20^. Recent evidence has further suggested that global cortical gray matter volume (GMV) may partially mediate the association between prenatal GDM and child adiposity outcomes in offspring^14^. These findings support the possibility that brain development as a critical intermediate phenotype linking prenatal metabolic dysregulation to later health outcomes.

However, current research has largely treated GDM exposure as a homogeneous developmental perturbation, typically focusing on average brain differences between GDM-exposed and unexposed youth^11–20^. This approach may obscure neuroanatomical heterogeneity within GDM-exposed offspring, whose clinical outcomes vary across anthropometric, neurocognitive, and psychiatric/behavioral domains. Because brain maturation follows coordinated and temporally ordered trajectories^21–23^, GDM-related neurodevelopmental differences may organize into distinct anatomical patterns rather than a single uniform alteration. Characterizing these patterns may improve our understanding of the heterogeneous clinical outcomes associated with prenatal GDM exposure.

In this study, we leveraged structural MRI (sMRI) data from the large ABCD® study to systematically characterize neurodevelopmental heterogeneity among GDM-exposed offspring. We applied Subtype and Stage Inference (SuStaIn), a data-driven approach, to identify distinct cortical thickness subtypes and infers their progression trajectory from cross-sectional data^24^. We then associated these subtypes with anthropometric, neurocognitive, psychiatric/behavioral, and anatomical measures across different time point. We hypothesized that prenatal exposure to GDM would be associated with heterogeneous cortical thickness patterns, rather than a single uniform pattern of anatomical alteration. Furthermore, we hypothesized that these subtype-specific patterns would be differentially associated with the aforementioned anthropometric, neurocognitive, psychiatric/behavioral, and anatomical outcomes.

## Methods

### Study sample

The ABCD® Study is an ongoing longitudinal study of adolescent development, with data collection at 21 study sites across the U.S. Data were obtained from the ABCD 5.1 release, including baseline, two-year, and four-year follow-up assessments. Participants were 9–10 years old at baseline and up to 15 years, 9 months old at the four-year follow-up. Information regarding funding agencies, recruitment sites, investigators, and project organizations can be obtained at https://abcdstudy.org. Details of the ABCD Study design have previously been reported^25,26^.

The primary sample included 573 youth exposed to GDM and 2,854 healthy controls (HCs). General exclusion criteria were consistent with our prior ABCD study and included major neurological, medical, developmental, and perinatal conditions^14,15^. Additional exclusions were applied to HCs with unhealthy weight status or past/current KSADS-defined neurodevelopmental or psychiatric diagnoses. Participants who failed neuroimaging quality control or had missing key covariates were also excluded.

### GDM exposure

GDM exposure was determined based on parental self-report at the baseline assessment, using this question: “During the pregnancy with this child, did you (or the biological mother) have pregnancy-related diabetes?” Responses were coded as a binary variable (yes/no).

### Outcome measures

#### Anthropometric measures

Youth’s weight (kg), height (cm), and waist circumference (cm) were measured by a trained researcher. Waist circumference was measured with a tape around the highest point on the pelvic bone. These measures were used to calculate waist-to-height ratio (WHtR) and BMI (kg/m^2^)^27^.

Age- and sex-specific BMI percentiles, BMI z-scores, weight z-scores, and height z-scores were calculated according to the Centers for Disease Control and Prevention (CDC) growth chart guidelines^28,29^. Childhood weight status was categorized based on CDC definitions: obesity (≥95th percentile), overweight (≥85th to <95th percentile), normal weight (≥5th to <85th percentile), and underweight (<5th percentile). Waist z-scores were derived from NHANES III reference data^30^.

#### Neurocognitive, psychiatric and behavioral assessment

General cognitive ability was evaluated using fully corrected T-scores from the NIH Toolbox Cognition Battery^31,32^. For psychiatric disorders, Kiddie Schedule for Affective Disorders and Schizophrenia (KSADS-5)^33^. The main analysis focused on current diagnosis of disorders, with past diagnoses reported in the Supplementary Materials. As a complementary dimensional measure, behavioral and emotional problems were assessed using T-scores from the Child Behavior Checklist (CBCL), a standardized parent-report inventory that yields syndrome scales for internalizing symptoms and externalizing behaviors^34^.

#### Image acquisition, processing and quality control

sMRI data were collected using standardized protocols across 21 ABCD study sites^35^. T1-weighted images were processed using FreeSurfer (version 5.3.0, https://surfer.nmr.mgh.harvard.edu/) for cortical surface reconstruction. The following measures were obtained: cortical gray and white matter volumes (mm^3^), regional cortical thickness (mm), and regional cortical surface area (mm^2^) based on Desikan-Killiany-Tourville (DKT) Atlas.

Neuroimaging analyses excluded participants who had abnormal radiological findings or whose T1 scan was of insufficient quality, as determined by the ABCD Data Analytics and Informatics Center^35^.Participants were also excluded based on quality control procedures on the cortical surface reconstruction for five categories of inaccuracy: severity of motion, intensity inhomogeneity, white matter underestimation, pial overestimation, and magnetic susceptibility artifact^35^.

### Data pre-processing

Imaging data were standardized and adjusted for relevant covariates. First, to mitigate inter-site variability, all sMRI measures were harmonized using ComBat for cross-sectional analyses and longitudinal ComBat harmonization for longitudinal analyses^36–38^, with site treated as a batch variable and age and sex included as biological covariates.

Following harmonization, sMRI measures were averaged across hemispheres. For the primary analysis, ROI-level measures were aggregated into six bilateral cortical regions—frontal, temporal, parietal, occipital, insula, and cingulate—for three reasons: 1) cortical thickness deviations associated with prenatal metabolic exposures are expected to be subtle during this developmental stage, and spatial aggregation may improve signal-to-noise; 2) reducing the number of features helps preserve statistical power given the available sample size; and 3) a lower-dimensional representation promotes model parsimony and stability. These aggregated ROIs refer to these collectively as “lobe-level” units, with the understanding that the insula and cingulate are not traditional neocortical lobes. Sensitivity analyses were additionally performed using ROI-level measures. Consistency between lobe- and ROI-level models was evaluated by comparing subtype assignments in the confusion matrix and progression ordering using Kendall’s τ after aggregating ROI-level events to the lobe level^39^.

SuStaIn model requires input features that follow a monotonic trajectory with respect to disease or exposure progression^24^. Therefore, we selected cortical thickness as the primary input feature because it peaks in very early childhood, around 2 years of age^40^, and is expected to decline during our observation window at ages 9–10 years. In contrast, surface area and subcortical volume generally peak later, around 12 years old, and therefore don’t satisfy the monotonic-decrease assumption within this age range. Importantly, inferred ordering patterns were interpreted as probabilistic differences in cortical organization rather than direct evidence of within-individual developmental progression.

To account for normal biological variation, all brain measures used as inputs to SuStaIn were residualized using the HC group as the reference. In addition, we regressed out the effects of age, sex, race/ethnicity, pubertal stage, parental education, household income, handedness, and estimated total intracranial volume (eTIV). Finally, all features were standardized to z-scores relative to the HC group to increase comparability across cortical lobes.

### SuStaIn modelling

SuStaIn is an unsupervised machine learning framework that identifies distinct subtypes based on shared phenotypic patterns across the continuum of disease progression or exposure-related effects. It infers stage-specific progression trajectories within each subtype and estimates the probability of subtype and stage for each individual based on cross-sectional data^24^. We further justified the applicability of SuStaIn to GDM-exposed youth in the Supplementary Materials. The input to SuStaIn was an M × N matrix, where M represents the number of participants (M = 573), and N denotes the number of biomarkers (N = 6), corresponding to cortical thickness across six lobes of interest. Given the relatively small deviations from HCs, we specified a single z-score threshold (z = 1) to define abnormality. The SuStaIn model was run using 25 random initializations and 100,000 Markov Chain Monte Carlo (MCMC) iterations to estimate the most probable sequence characterizing the spatiotemporal progression of cortical changes.

First, we used the Hopkins statistic to assess whether the data exhibited a non-random clustering tendency^41^. SuStaIn models were then fitted sequentially with varying numbers of subtypes (K = 1–3). The optimal number of subtypes was determined using five-fold cross-validation by selecting the model with the lowest Cross-Validation Information Criterion (CVIC)^42^. Second, to evaluate the stability of subtype progression patterns across folds, we computed the Cross-Validation Similarity (CVS), defined as the average Bhattacharyya coefficient between pairs of subtype progression sequences^43^. Finally, after identifying the optimal K, everyone was assigned to the most likely subtype and disease stage based on the maximum-likelihood estimates derived from the MCMC samples.

## Statistical analysis

After SuStaIn modeling, each child was assigned to the subtype with the highest posterior likelihood based on the MCMC samples. Baseline offspring characteristics were compared across five domains: demographic, anthropometric, neurocognitive, psychiatric/behavioral, and sMRI measures. Continuous variables with approximately normal distributions were compared using two-sample t-tests, whereas categorical variables were compared using chi-square or Fisher’s exact tests, as appropriate.

Longitudinal associations were examined using linear mixed-effects models with subtype, time, and their interaction as predictors across available baseline, 2-year, and 4-year follow-up data. Statistical significance was defined as p < 0.05, with false discovery rate (FDR) correction applied for multiple comparisons. The full analytic pipeline is illustrated in Figure 1.

**Figure 1.**
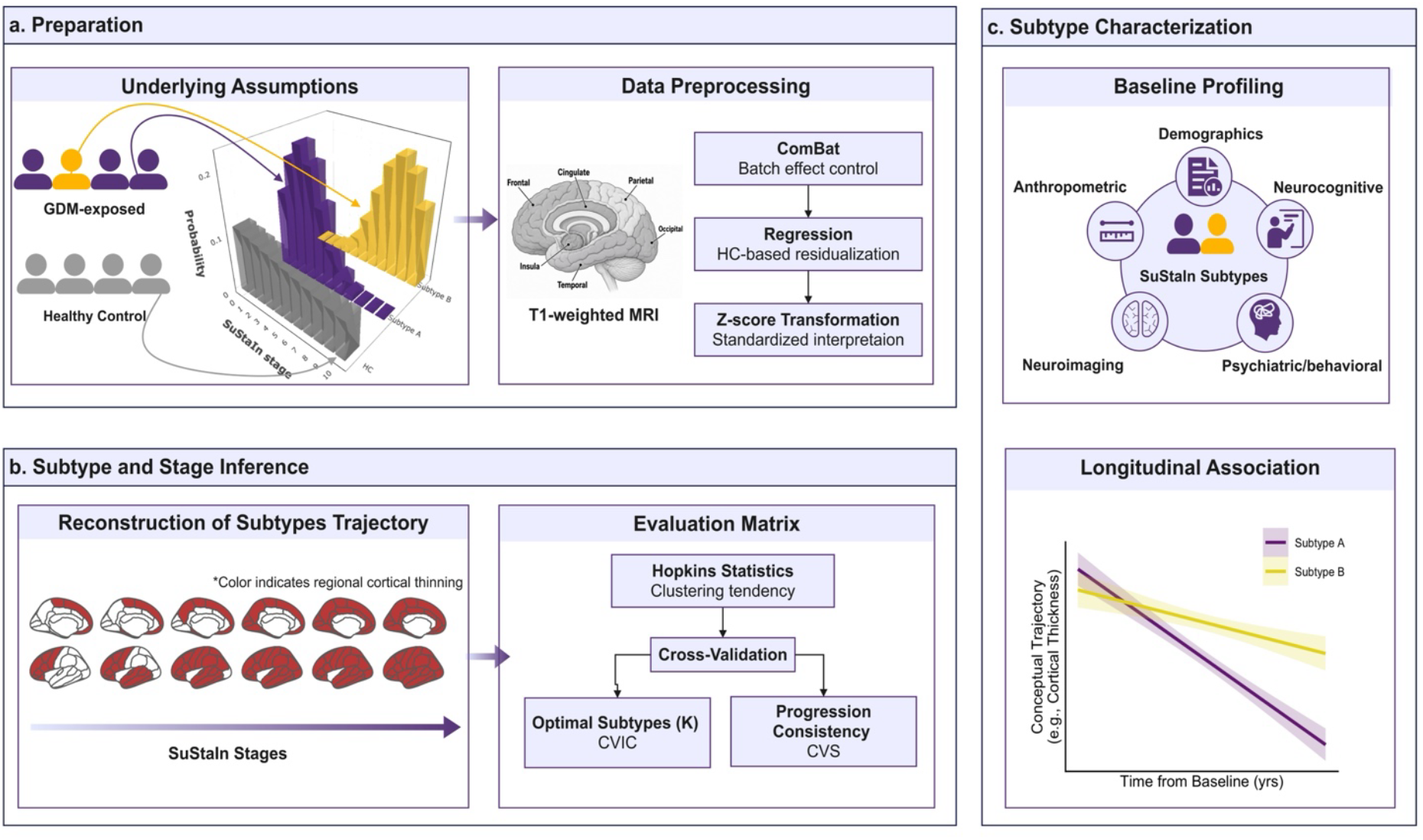
Analytical framework for identifying neuroanatomical heterogeneity in GDM-exposed children. (a) Preparation and preprocessing: The histogram illustrates a schematic subtype–stage probability distribution under the SuStaIn framework. Structural T1-weighted MRI data from GDM-exposed children and demographically matched healthy controls were harmonized using ComBat, residualized based on HC regression models, and standardized via z-score transformation. (b) Subtype and stage inference (SuStaIn): The probabilistic SuStaIn model was applied to infer latent subtypes and their stage-wise progression patterns. The evaluation matrix included the Hopkins statistic to assess clustering tendency, cross-validation for model stability, the cross-validation information criterion (CVIC) to determine the optimal number of subtypes, and the cross-validation similarity (CVS) to quantify progression consistency. (c) Subtype-level characterization and longitudinal association: Identified subtypes were profiled at baseline across demographic, anthropometric, pregnancy-related, neuroimaging, cognitive, and psychiatric domains. Linear mixed-effects models evaluated longitudinal trajectories to determine whether subtype membership was associated with differential developmental patterns over time.

## Data and code availability

The data that support the findings of this study are available from the ABCD. Restrictions apply to the availability of these data, which were used under license for this study. Data are available from the authors with the permission of ABCD. SuStaIn algorithm is available on the UCL-POND GitHub (https://github.com/ucl-pond/).

## Results

Table 1 displays characteristics of our study sample. Among the GDM-exposed youth (*N* = 573), the mean age was 119.02 months (SD = 7.34), and the sample was nearly evenly split by sex (52.5% male, 47.5% female). Most participants were White (59.7%), and in healthy weight status (55.7%). Baseline comparisons between GDM-exposed youth and HCs are presented in Supplementary Table 1.

**Table 1.** Demographic, anthropometric, and psychiatric characteristics by Subtypes.

| Variable | Subtype 1<br>(N=362) | Subtype 2<br>(N=211) | Total<br>(N=573) | Effect<br>Size | p value | pFDR |
| --- | --- | --- | --- | --- | --- | --- |
| **Age, months** | 119.51 (7.46) | 118.18 (7.08) | 119.02 (7.34) | -0.18 | **0.036** | 0.216 |
| **Sex (Female)** | 169 (46.7%) | 103 (48.8%) | 272 (47.5%) | 0 | 0.622 | 0.963 |
| **Race/ethnicity** |  |  |  | 0 | 0.644 | 0.963 |
| White | 211 (58.3%) | 131 (62.1%) | 342 (59.7%) |  |  |  |
| Black | 50 (13.8%) | 28 (13.3%) | 78 (13.6%) |  |  |  |
| Others | 101 (27.9%) | 52 (24.6%) | 153 (26.7%) |  |  |  |
| **Family income** |  |  |  | 0.02 | 0.329 | 0.963 |
| [<50K] | 126 (34.8%) | 61 (28.9%) | 187 (32.6%) |  |  |  |
| [>=100K] | 130 (35.9%) | 80 (37.9%) | 210 (36.6%) |  |  |  |
| [>=50K & <100K] | 106 (29.3%) | 70 (33.2%) | 176 (30.7%) |  |  |  |
| **Parental education (bachelor's degree or above)** | 199 (55.0%) | 117 (55.5%) | 316 (55.1%) | 0 | 0.912 | 0.963 |
| **Pubertal stage** |  |  |  | 0 | 0.963 | 0.963 |
| Prepuberty | 169 (46.7%) | 101 (47.9%) | 270 (47.1%) |  |  |  |
| Early puberty | 88 (24.3%) | 50 (23.7%) | 138 (24.1%) |  |  |  |
| Mid-post puberty | 105 (29.0%) | 60 (28.4%) | 165 (28.8%) |  |  |  |
| **Weight status** |  |  |  | 0.06 | 0.187 | 0.28 |
| Underweight | 12 (3.3%) | 4 (1.9%) | 16 (2.8%) |  |  |  |
| Healthy | 190 (52.5%) | 129 (61.1%) | 319 (55.7%) |  |  |  |
| Overweight | 67 (18.5%) | 36 (17.1%) | 103 (18.0%) |  |  |  |
| Obese | 93 (25.7%) | 42 (19.9%) | 135 (23.6%) |  |  |  |
| **BMI** | 19.99 (4.93) | 19.24 (4.27) | 19.71 (4.71) | -0.16 | 0.064 | 0.129 |
| **BMI-z** | 0.68 (1.27) | 0.59 (1.13) | 0.65 (1.22) | -0.07 | 0.398 | 0.437 |
| **WHtR** | 0.49 (0.08) | 0.49 (0.07) | 0.49 (0.07) | -0.07 | 0.437 | 0.437 |
| **Height(cm)** | 142.03 (8.25) | 139.20 (7.06) | 140.99 (7.95) | -0.36 | **< 0.001** | **< 0.001** |
| **Weight(kg)** | 40.79 (12.46) | 37.66 (10.70) | 39.64 (11.93) | -0.26 | **0.002** | **0.007** |
| **ADHD<sup>a</sup>** | 17 (4.7%) | 19 (9.2%) | 36 (6.4%) | 0.08 | **0.036** | 0.249 |
| **Disruptive Behavior Disorder<sup>a</sup>** | 13 (3.6%) | 12 (5.8%) | 25 (4.4%) | 0.03 | 0.23 | 0.599 |
| **Anxiety<sup>a</sup>** | 40 (11.1%) | 26 (12.5%) | 66 (11.6%) | 0 | 0.627 | 0.732 |
| **OCD<sup>a</sup>** | 30 (8.4%) | 12 (5.8%) | 42 (7.4%) | 0.02 | 0.257 | 0.599 |
| **MDD/Persistent Depressive Disorder<sup>a</sup>** | 0 (0.0%) | 0 (0.0%) | 0 (0.0%) |  |  |  |
| **Bipolar Disorder<sup>a</sup>** | 3 (0.8%) | 2 (1.0%) | 5 (0.9%) | 0 | 0.877 | 0.877 |
| **Feeding/Eating Disorder<sup>a</sup>** | 3 (0.8%) | 3 (1.4%) | 6 (1.1%) | 0 | 0.496 | 0.695 |
| **PTSD<sup>a</sup>** | 1 (0.3%) | 0 (0.0%) | 1 (0.2%) | 0 | 0.446 | 0.695 |

**Notes:** Values are presented as *n* (%) for categorical variables and *mean* (*SD*) for continuous variables. Effect sizes are reported as Cohen's *d* for continuous variables (reference on Subtype 1) and Cramér's *V* for categorical variables. Effect sizes are rounded to two decimal places and values displayed as 0 indicate very small (non-zero) effects. P values were obtained using two-sample t-tests for continuous variables and chi-square or Fisher's exact tests for categorical variables, as appropriate. P values ≤ 0.05 are shown in bold; FDR correction was applied separately within demographic, anthropometric, and psychiatric variable groups.
BMI = body mass index; BMI-z = BMI z-score; WHtR = waist-to-height ratio. ADHD = attention-deficit/hyperactivity disorder; OCD = obsessive-compulsive disorder; MDD = major depressive disorder; BD = bipolar disorder; PTSD = posttraumatic stress disorder.
a. Sample size for these analyses: *n* = 567 (Subtype 1: *n* = 322; Subtype 2: *n* = 245).

### Subtype patterns

The Hopkins statistic of the input features was 0.76 (95% CI: 0.74 to 0.76), indicating a moderate clustering tendency. SuStaIn was then applied to baseline cortical thickness data to infer subtype-specific patterns. The optimal number of subtypes was determined as two based on the lowest Cross-Validation Information Criterion (CVIC = 10,998.33; Supplementary Figure 1).

Figure 2a illustrates the inferred stage-wise ordering of cortical thinning events for each subtype. In Subtype 1, the highest-probability sequence suggested early involvement of the insula, followed by cingulate, occipital and parietal regions, with later extension to the frontal and temporal. In contrast, Subtype 2 demonstrated an alternative probabilistic ordering pattern, beginning in the frontal lobe and subsequently involving the temporal, parietal, occipital, insula, and finally cingulate regions. These sequences represent subtype-specific differences in the relative ordering of regional abnormalities inferred at the group level.

**Figure 2.**
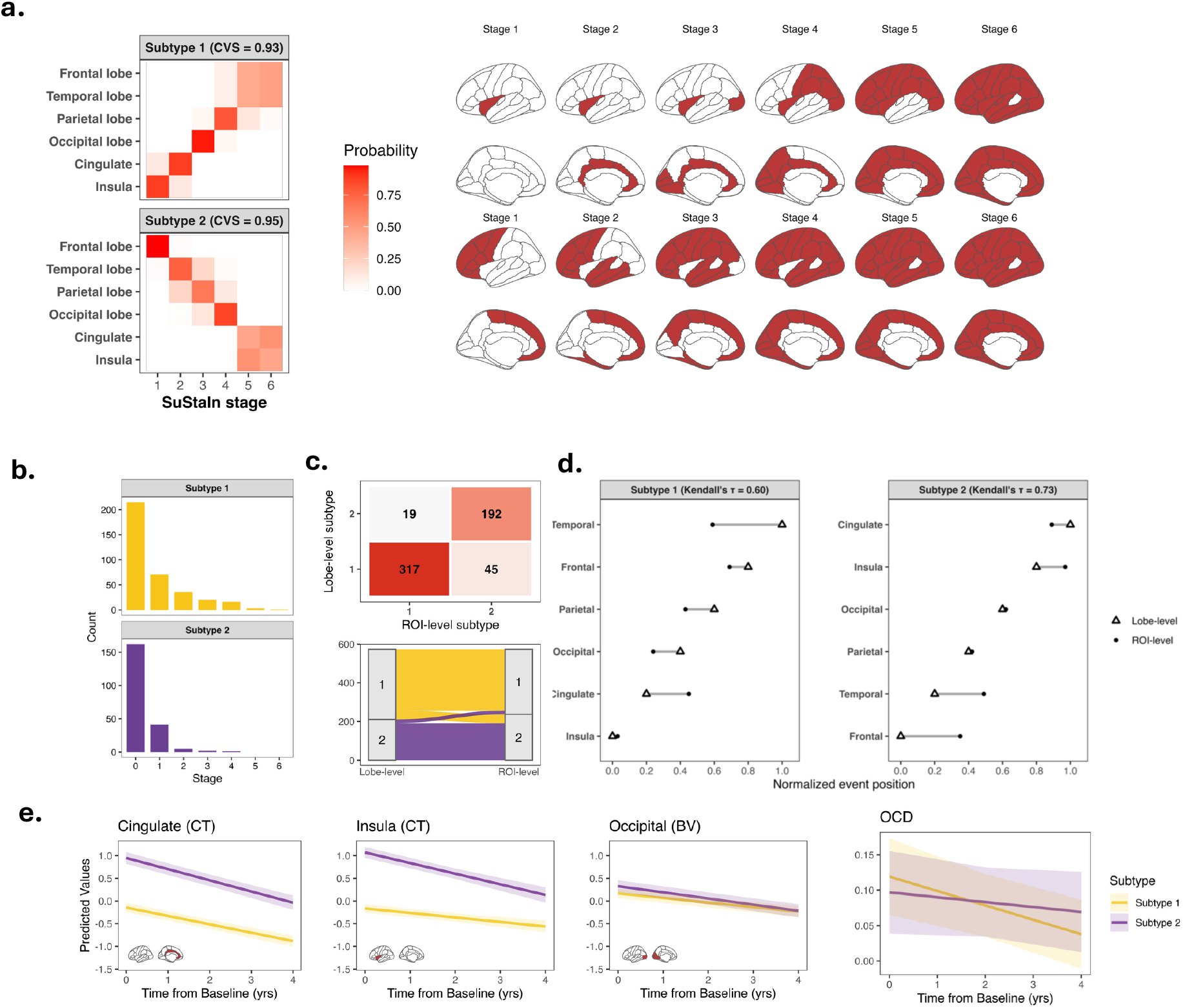
Identification and validation of cortical thinning subtypes among children with prenatal GDM exposure. **(a) Spatiotemporal progression patterns.** The optimal number of subtypes was determined to be two based on the lowest CVIC. For each subtype, the left heatmap displays the probability that the z-score of each cortical lobe reaches the atrophy threshold at each SuStaIn stage, with warmer colors indicating a higher likelihood of earlier involvement. High CVS values indicate high progression stability. The right panels show reconstructed brain surface maps illustrating the sequential spatial accumulation of cortical atrophy across six SuStaIn stages for each subtype, highlighting distinct anatomical progression patterns. **(b) Stage distribution by subtypes**. Stage distributions show the inferred disease stage of individuals within each subtype. **(c) Consistency of subtype assignments across lobe-level and ROI-level models**. Confusion matrix (top) and alluvial plot (bottom) compare subtype assignments derived from the lobe-level model and the ROI-level model. Most participants retained the same subtype classification across models, indicating strong agreement between feature resolutions. **(d) Comparison of event ordering between lobe-level and ROI-level models**. Normalized event positions for each cortical region are shown for Subtype 1 and Subtype 2. Triangles indicate the event ordering inferred from the lobe-level model, and dots indicate the ordering derived from the ROI-level model. The ordering showed substantial concordance between models (Subtype 1: Kendall’s τ = 0.60; Subtype 2: Kendall’s τ = 0. 73), supporting the robustness of subtype progression patterns. **(e) Longitudinal subtype differences in neuroanatomical and psychiatric outcomes**. Lines represent model-predicted values from LMM. Shaded bands indicate 95% confidence intervals. CT= Cortical Thickness; BV = Brain Volume.

To assess the robustness of the subtyping, we repeated the SuStaIn model using ROI-level cortical thickness features as a sensitivity analysis. Subtype assignments were largely consistent between the two models, with 88.9 % of the participants retaining the same subtype (Figure 2b–c). We further compared the inferred progression ordering by aggregating ROI-level results to the lobe level. The event sequences showed moderate to high concordance across models (Subtype 1: Kendall’s τ = 0.60; Subtype 2: τ = 0.73; Figure 2d).

Despite minor shifts in the relative staging of several lobes, the overall progression structure was preserved, supporting the robustness of the subtype identified in the primary lobe-level model.

### Subtype assignments and phenotypic differences

Among GDM-exposed youth, 362 (63.2%) were classified as Subtype 1 and 211 (36.8%) were classified as Subtype 2. Subtype 1 was slightly older than Subtype 2 (119.51 ± 7.46 vs. 118.18 ± 7.08 months; *d* = 0.18, *p* = 0.036, *p*_*FDR*_ = 0.216). No significant differences were observed in sex distribution, race/ethnicity, household income, parental education or pubertal stage.

For anthropometric measures, Subtype 1 had greater height and weight than Subtype 2 (142.03 ± 8.25 vs. 139.20 ± 7.06 cm; 40.79 ± 12.46 vs. 37.66 ± 10.70 kg), with small-to-moderate effect sizes (height: *d* = 0.36, *p*_*FDR*_ < 0.001; weight: *d* = 0.26, *p*_*FDR*_ = 0.007). Weight status, BMI, BMI-z and WHtR did not significantly differ between groups.

For neurocognitive, behavioral and psychiatric measures, several nominal group differences were observed. Subtype 2 had a higher prevalence of ADHD diagnosis than Subtype 1 (9.2% vs. 4.7%; *d* = 0.08, *p* = 0.036, *p*_*FDR*_ = 0.249), consistent with higher CBCL ADHD T-scores in Subtype 2 (53.61 ± 6.31 vs. 52.64 ± 4.86; *d* = 0.18, *p* = 0.040, *p*_*FDR*_ = 0.533). Subtype 2 also showed a higher prevalence of past bipolar disorder than Subtype 1 (11.1% vs. 5.3%; *d* = 0.10, *p* = 0.012, *p*_*FDR*_ = 0.092). However, these effects didn’t survive FDR correction. No significant differences emerged in other KSADS domains, NIH Toolbox cognitive measures, or other CBCL scores (Supplementary Table 2, 3, 4).

For anatomical measures, significant differences were observed between Subtype 1 and Subtype 2 (Supplementary Table 5). Compared with Subtype 1, Subtype 2 exhibited greater cortical thickness across all lobes (all *p*_*FDRs*_ < 0.001), with the largest effect observed in the insula (*d* = 1.64). Subtype 2 also showed larger volumes, although the effect sizes were generally small, with the largest effect sizes observed in the temporal lobe (*d* = 0.32, *p*_*FDR*_ = 0.002), insula (*d* = 0.24, *p*_*FDR*_ = 0.018) and parietal lobe (*d* = 0.23, *p* = 0.018). In contrast, Subtype 1 demonstrated significantly larger surface area across all lobes, with the largest difference observed in the insula (*d* = 0.31, *p*_*FDR*_ *= 0*.*002*), frontal (*d* = 0.29, *p*_*FDR*_ = 0.003), cingulate (*d* = -0.27, *p*_*FDR*_ = 0.002) and occipital (*d* = 0.19, *p* =0.004) lobes.

### Longitudinal trajectory

We examined longitudinal associations by modeling subtype, time (years since baseline), and their interaction on standardized cortical thickness, volume, and surface area across six lobes (Table 2). For cortical thickness, significant interactions were observed across all six lobes (β range: −0.05 to −0.13, all *p* < 0.05, all *p*_*FDRs*_ < 0.05, Figure 2e). These consistently negative interaction effects indicate that Subtype 2 showed a steeper longitudinal decline in cortical thickness compared with Subtype 1, with the largest effect observed in the insula (β = -0.13, 95% CI: −0.19 to −0.08, *p*_*FDR*_ < 0.001).

**Table 2.** Longitudinal Effects of Subtype and Time on sMRI (observations =1146).

| Terms | Cortical Thickness |  |  |  | Brain Volume |  |  |  | Surface Area |  |  |  |
| --- | --- | --- | --- | --- | --- | --- | --- | --- | --- | --- | --- | --- |
| | $\beta$ | 95% CI | p | pFDR | $\beta$ | 95% CI | p | pFDR | $\beta$ | 95% CI | p | pFDR |
| **Frontal** |  |  |  |  |  |  |  |  |  |  |  |  |
| subtype2 | 1.22 | (1.10, 1.35) | **<0.001** | **<0.001** | 0.21 | (0.10, 0.31) | **<0.001** | **<0.001** | -0.28 | (-0.39, -0.17) | **<0.001** | **<0.001** |
| time | -0.23 | (-0.26, -0.20) | **<0.001** | **<0.001** | -0.14 | (-0.16, -0.13) | **<0.001** | **<0.001** | 0.07 | (-0.00, 0.14) | 0.066 | 0.192 |
| interaction | -0.05 | (-0.10, -0.00) | **0.037** | **0.037** | -0.02 | (-0.04, 0.01) | 0.201 | 0.201 | 0.01 | (-0.01, 0.03) | 0.342 | 0.683 |
| **Parietal** |  |  |  |  |  |  |  |  |  |  |  |  |
| subtype2 | 1 | (0.86, 1.13) | **<0.001** | **<0.001** | 0.22 | (0.10, 0.34) | **<0.001** | **<0.001** | -0.21 | (-0.34, -0.08) | **0.002** | **0.002** |
| time | -0.23 | (-0.26, -0.20) | **<0.001** | **<0.001** | -0.19 | (-0.21, -0.18) | **<0.001** | **<0.001** | 0.06 | (-0.02, 0.13) | 0.128 | 0.192 |
| interaction | -0.05 | (-0.10, -0.00) | **0.033** | **0.037** | -0.02 | (-0.04, 0.00) | 0.11 | 0.164 | 0.02 | (-0.00, 0.04) | 0.071 | 0.428 |
| **Occipital** |  |  |  |  |  |  |  |  |  |  |  |  |
| subtype2 | 0.71 | (0.55, 0.87) | **<0.001** | **<0.001** | 0.16 | (0.02, 0.30) | **0.029** | **0.035** | -0.24 | (-0.39, -0.10) | **<0.001** | **0.002** |
| time | -0.16 | (-0.19, -0.13) | **<0.001** | **<0.001** | -0.1 | (-0.12, -0.08) | **<0.001** | **<0.001** | 0.06 | (-0.02, 0.13) | 0.121 | 0.192 |
| interaction | -0.06 | (-0.11, -0.01) | **0.032** | **0.037** | -0.04 | (-0.06, -0.01) | **0.004** | **0.026** | -0.01 | (-0.02, 0.01) | 0.489 | 0.734 |
| **Temporal** |  |  |  |  |  |  |  |  |  |  |  |  |
| subtype2 | 1.18 | (1.03, 1.32) | **<0.001** | **<0.001** | 0.32 | (0.21, 0.43) | **<0.001** | **<0.001** | -0.18 | (-0.30, -0.07) | **0.002** | **0.002** |
| time | -0.14 | (-0.17, -0.11) | **<0.001** | **<0.001** | -0.12 | (-0.14, -0.11) | **<0.001** | **<0.001** | 0.09 | (0.02, 0.16) | 0.015 | 0.092 |
| interaction | -0.07 | (-0.12, -0.01) | **0.014** | **0.027** | -0.02 | (-0.04, 0.01) | 0.183 | 0.201 | 0.01 | (-0.01, 0.03) | 0.274 | 0.683 |
| **Cingulate** |  |  |  |  |  |  |  |  |  |  |  |  |
| subtype2 | 1.09 | (0.95, 1.24) | **<0.001** | **<0.001** | 0.12 | (-0.01, 0.26) | 0.077 | 0.077 | -0.27 | (-0.41, -0.13) | **<0.001** | **<0.001** |
| time | -0.18 | (-0.21, -0.16) | **<0.001** | **<0.001** | -0.1 | (-0.12, -0.09) | **<0.001** | **<0.001** | 0 | (-0.08, 0.09) | 0.947 | 0.947 |
| interaction | -0.06 | (-0.10, -0.02) | **0.004** | **0.012** | -0.02 | (-0.04, 0.00) | 0.078 | 0.156 | 0 | (-0.02, 0.03) | 0.959 | 0.959 |
| **Insula** |  |  |  |  |  |  |  |  |  |  |  |  |
| subtype2 | 1.23 | (1.10, 1.37) | **<0.001** | **<0.001** | 0.26 | (0.12, 0.39) | **<0.001** | **<0.001** | -0.3 | (-0.44, -0.16) | **<0.001** | **<0.001** |
| time | -0.1 | (-0.13, -0.07) | **<0.001** | **<0.001** | -0.03 | (-0.06, -0.01) | **0.011** | **<0.001** | 0.01 | (-0.09, 0.10) | 0.903 | 0.947 |
| interaction | -0.13 | (-0.19, -0.08) | **<0.001** | **<0.001** | -0.05 | (-0.09, -0.01) | **0.011** | **0.033** | 0.01 | (-0.03, 0.06) | 0.647 | 0.776 |

**Notes:** Results from linear mixed-effects models assessing subtype, time, and their interaction on cortical thickness (after z-score standardization), volume, and surface area across brain regions. Cortical thickness, brain volume, and surface area were adjusted for sex and baseline age. Brain volume and surface area were additionally adjusted for estimated total intracranial volume (eTIV); Bolded values are statistically significant, p stands for raw p values and pFDR stands for p values after FDR correction.

For brain volume, significant subtype-by-time interactions in the occipital lobe (β = −0.04, 95% CI:−0.06 to −0.01, *p*_*FDR*_ = 0.026) and insula (β = −0.05, 95% CI: −0.09 to −0.01, *p*_*FDR*_ = 0.033) were observed. These interaction effects indicate that Subtype 2 exhibited a steeper longitudinal decline in specific brain volume regions compared with Subtype 1. No significant subtype × time interactions were detected for surface area.

We further examined longitudinal associations of subtype, time, and their interaction with anthropometric and psychiatric outcomes (Table 3). For anthropometric measures, no subtype-by-time interaction survived FDR correction. For psychiatric outcomes, the subtype-by-time interaction for OCD symptoms remained significant (β = 0.91, 95% CI: 0.24 to 1.59, *p* = 0.008, *p*_*FDR*_ = 0.049). This significant interaction indicates a slower decline in OCD prevalence over time in Subtype 2. Other psychiatric outcomes did not show significant subtype-by-time interactions after FDR correction. Longitudinal results for CBCL and NIH Toolbox outcomes are provided in Supplementary Tables 6 and 7, no significance in interaction was observed.

**Table 3.** Longitudinal associations of subtype and time with anthropometric (observations = 1853) and psychiatric outcomes (observations = 1331).

| Outcome | Terms | Estimate | 95% CI | p | pFDR |
| --- | --- | --- | --- | --- | --- |
| **BMI** | subtype2 | -0.58 | (-1.41, 0.25) | 0.175 | 0.175 |
|  | time | 0.94 | (0.82, 1.06) | **0.001** | **<0.001** |
|  | interaction | 0.07 | (-0.07, 0.22) | 0.331 | 0.331 |
| **Height(cm)** | subtype2 | -2.33 | (-3.45, -1.21) | **0.001** | **<0.001** |
|  | time | 5.43 | (5.21, 5.64) | **0.001** | **<0.001** |
|  | interaction | -0.26 | (-0.52, -0.00) | **0.049** | 0.146 |
| **Weight(kg)** | subtype2 | -2.55 | (-4.82, -0.29) | **0.028** | **0.041** |
|  | time | 5.8 | (5.45, 6.16) | **0.001** | **<0.001** |
|  | interaction | -0.22 | (-0.65, 0.21) | 0.321 | 0.331 |
| **ADHD** | subtype2 | 0.97 | (-0.28, 2.23) | 0.129 | 0.388 |
|  | time | 0.02 | (-0.40, 0.45) | 0.917 | 0.917 |
|  | interaction | 0.09 | (-0.36, 0.54) | 0.686 | 0.946 |
| **Disruptive Behavior Disorder** | subtype2 | -0.17 | (-1.45, 1.11) | 0.795 | 0.795 |
|  | time | 0.08 | (-0.31, 0.47) | 0.699 | 0.839 |
|  | interaction | -0.07 | (-0.55, 0.42) | 0.788 | 0.946 |
| **Anxiety** | subtype2 | -0.10 | (-0.83, 0.63) | 0.793 | 0.795 |
|  | time | -0.27 | (-0.53, -0.00) | **0.047** | 0.140 |
|  | interaction | 0.11 | (-0.22, 0.43) | 0.518 | 0.946 |
| **OCD** | subtype2 | -1.22 | (-2.77, 0.33) | 0.122 | 0.388 |
|  | time | -1.02 | (-1.61, -0.44) | **0.001** | **0.004** |
|  | interaction | 0.91 | (0.24, 1.59) | **0.008** | **0.049** |
| **BD** | subtype2 | 0.28 | (-1.40, 1.95) | 0.745 | 0.795 |
|  | time | -0.67 | (-1.74, 0.41) | 0.227 | 0.453 |
|  | interaction | 0.37 | (-0.87, 1.61) | 0.558 | 0.946 |
| **Feeding/Eating Disorder** | subtype2 | 0.67 | (-0.96, 2.29) | 0.422 | 0.795 |
|  | time | -0.26 | (-1.04, 0.52) | 0.509 | 0.764 |
|  | interaction | -0.02 | (-1.02, 0.98) | 0.968 | 0.968 |

**Notes:** Results from regression models assessing subtype, time, and their interaction on anthropometric (BMI, height, weight) and KSADS psychiatric outcomes. MDD and PTSD were excluded from the GLMM analyses because low prevalence resulted in insufficient case counts for reliable model estimation. Continuous outcomes were analyzed using linear mixed-effects models, and psychiatric outcomes were analyzed using generalized linear mixed-effects models (GLMMs). For consistency, estimates from GLMMs are presented on the linear scale rather than as odds ratios. All models were adjusted for baseline age, sex, race, household income, parental education, and puberty stage. Bolded values are statistically significant, p stands for raw p value and pFDR stands for p values after FDR correction. ADHD = attention-deficit/hyperactivity disorder; OCD = obsessive-compulsive disorder; BD = bipolar disorder.

## Discussion

In this study, we applied a data-driven SuStaIn model to characterize anatomical heterogeneity of GDM-exposed youth. We identified two distinct cortical thickness patterns. Subtype 1 was characterized by earlier insular-cingulate regions thinning and was associated with greater height and weight. In contrast, Subtype 2 was characterized by earlier inferred frontal involvement, steeper longitudinal cortical thinning, and nominally greater psychiatric-related measures.

Subtype 1 exhibited a distinct anatomical sequence initiated by early thinning in the insula and cingulate cortices. This subtype was associated with a significant somatic divergence at baseline, with these individuals displaying greater height and weight that persisted, but did not accelerate, across the longitudinal follow-up. The insula is the primary brain center for interoceptive processing, sensing internal metabolic and satiety signals^44–46^, while the anterior cingulate plays a pivotal role in the top-down regulation of appetitive behaviors^47–49^. By reaching a fixed developmental state prematurely prior to late childhood, these regulatory hubs may have established a higher metabolic set-point programmed in response to the intrauterine GDM environment. This is manifested in our cohort as the stable, greater weight and height growth trajectory observed in Subtype 1.

Subtype 2 demonstrated relatively earlier inferred frontal and temporal lobes, a subtype that aligns with the high baseline prevalence of ADHD in this group. Unlike Subtype 1, Subtype 2 demonstrates a psychiatric escalation, evidenced by a longitudinal increase in the odds of OCD diagnosis. These disorders share impairments in executive function, impulse control, and behavioral regulation, domains supported by prefrontal and frontostriatal circuits, and often associated with accelerated frontal thinning^50–52^. The subsequent spread of thinning to parietal and occipital regions may reflect a broader disruption of association networks following early frontal involvement. Notably, this pattern appears partially independent of anthropometric measures, suggesting that GDM-exposed youth with healthy weight status may still exhibit increased neurodevelopmental and psychiatric vulnerability.

Cross-sectional anatomical analysis provided further insight into these divergent trajectories. At baseline, Subtype 2 exhibited greater cortical thickness and larger brain volumes, but significantly smaller cortical surface area compared to Subtype 1. Because cortical thickness and surface area reflect partially independent neurodevelopmental processes^53–55^, this pattern may indicate differences in the balance between cortical expansion and synaptic pruning. The thicker cortex observed in Subtype 2 may reflect relatively delayed cortical thinning, whereas the larger surface area in Subtype 1 may suggest greater cortical expansion and a more advanced stage of cortical maturation during this developmental window.

Our longitudinal analysis of sMRI provides critical validation of these cross-sectional subtypes. While cortical thinning is a normative process during adolescence, reflecting synaptic pruning^22,56^, Subtype 2 exhibited a significantly steeper age-related thinning. This accelerated thinning corroborates the “frontal-first” progression identified by the SuStaIn model, which may reflect accelerated synaptic pruning or premature anatomical maturation, a process frequently observed in pediatric psychiatric disorders^57^. This “accelerated thinning” of the cortex may be the mechanistic driver linking prenatal GDM exposure to adolescent psychiatric onset. For example, GDM-induced oxidative stress may accelerate the cellular clock of frontal lobe maturation, reducing synaptic plasticity and precipitating the psychiatric vulnerabilities observed in this subgroup^58^.

Because subtype assignments were derived in part from cortical thickness features, cross-sectional differences in cortical thickness between subtypes should be interpreted cautiously, as they may partially reflect the model structure. Nevertheless, the accompanying differences in surface area, brain volume, longitudinal cortical thinning, and clinical outcomes suggest that the identified subtypes may capture broader neurodevelopmental variation beyond baseline cortical thickness alone.

### Limitation and Future Direction

Several limitations should be acknowledged. First, as the SuStaIn model was fit to cross-sectional baseline data, the inferred ordering represents a probabilistic model of between-subject heterogeneity rather than direct evidence of within-person developmental progression. In addition, SuStaIn assumes a monotonic pattern of biomarker progression, which may not fully capture nonlinear or potentially bidirectional neurodevelopmental changes during childhood and adolescence. Although longitudinal analyses offer supporting evidence for subtype-related differences in cortical change, definitive characterization of developmental trajectories and later-emerging clinical outcomes will require longer-term follow-up with denser repeated measurements. Future studies using longitudinal latent growth models, normative modeling frameworks, or recurrent deep learning approaches that do not require monotonic trajectories may better characterize complex neurodevelopmental changes associated with prenatal GDM exposure. Second,between-subtype differences in psychiatric and anthropometric measures were modest, albeit statistically significant, and may be amplified in larger samples with broader age ranges. While the model demonstrated stability across cross-validation folds and consistency between lobe-level and ROI-level analyses, further validation in independent cohorts is required to establish the generalizability and robustness of these subtypes. Third, GDM status relied on retrospective caregiver reports without granular data on disease severity or glycemic control, such as maternal HbA1c, limiting our ability to test dose-dependent associations with anatomical severity. Finally, disentangling prenatal programming from the postnatal environment remains challenging; unmeasured factors such as childhood lifestyle factors may compound prenatal risks, potentially influencing the observed anatomical trajectories.

## Conclusion

In summary, our findings suggest that prenatal exposure to GDM may be associated with heterogeneous neurodevelopmental trajectories during peri-adolescence. By applying a data-driven progression model SuStaIn, we identified two divergent developmental pathways: a ‘somatic-metabolic’ phenotype driven by insula-limbic dysregulation and systemic anabolic overgrowth, and a ‘ neuropsychiatric’ phenotype characterized by accelerated frontal lobe thinning and vulnerability to psychiatric disorders. These findings underscore the critical importance of moving beyond a ‘one-size-fits-all’ model of risk. Effective clinical surveillance must recognize these distinct trajectories, differentiating between youth who primarily require metabolic monitoring and those warranting early psychiatric intervention, ultimately paving the way for more precise, biologically informed strategies to mitigate the adverse consequences of GDM in offspring.

## Acknowledgement

The authors acknowledge and give thanks to the participants of the ABCD Study and their families.

Research described in this study was supported by the National Institutes of Health (NIH) R01DK137899 (PI: S.L.), the Brain and Behavior Research Foundation NARSAD Young Investigator award, the NIH K01MH122774, and NIH R01MH133840 (PI: Z.X.). Data used in the preparation of this article were obtained from the Adolescent Brain Cognitive Development (ABCD) Study (https://abcdstudy.org), held in the NIMH Data Archive (NDA). This is a multisite, longitudinal study designed to recruit more than 10,000 youth aged 9– 10 and follow them over 10 years into early adulthood. The ABCD Study is supported by the National Institutes of Health and additional federal partners under award numbers U01DA041048, U01DA050989, U01DA051016, U01DA041022, U01DA051018, U01DA051037, U01DA050987, U01DA041174, U01DA041106, U01DA041117, U01DA041028, U01DA041134, U01DA050988, U01DA051039, U01DA041156, U01DA041025, U01DA041120, U01DA051038, U01DA041148, U01DA041093, U01DA041089, U24DA041123, U24DA041147. A full list of supporters is available at https://abcdstudy.org/federal-partners.html. A listing of participating sites and a complete listing of the study investigators can be found at https://abcdstudy.org/consortium_members/. ABCD consortium investigators designed and implemented the study and/or provided data but did not necessarily participate in the analysis or writing of this report. This manuscript reflects the views of the authors and may not reflect the opinions or views of the NIH or ABCD consortium investigators.

## Reference

1. Boney CM, Verma A, Tucker R, Vohr BR. Metabolic syndrome in childhood: Association with birth weight, maternal obesity, and gestational diabetes mellitus. Pediatrics. 2005;115(3):e290–e296. doi:10.1542/peds.2004-1808

2. Catalano PM, McIntyre HD, Cruickshank JK, et al. The hyperglycemia and adverse pregnancy outcome study: Associations of GDM and obesity with pregnancy outcomes. Diabetes Care. 2012;35(4):780–786. doi:10.2337/dc11-1790

3. Kubo A, Ferrara A, Windham GC, et al. Maternal hyperglycemia during pregnancy predicts adiposity of the offspring. Diabetes Care. 2014;37(11):2996–3002. doi:10.2337/dc14-1438

4. Page KA, Romero A, Buchanan TA, Xiang AH. Gestational diabetes mellitus, maternal obesity, and adiposity in offspring. J Pediatr. 2014;164(4):807–810. doi:10.1016/j.jpeds.2013.11.063

5. Nahum Sacks K, Friger M, Shoham-Vardi I, et al. Prenatal exposure to gestational diabetes mellitus as an independent risk factor for long-term neuropsychiatric morbidity of the offspring. Am J Obstet Gynecol. 2016;215(3):380.e1-380.e7. doi:10.1016/j.ajog.2016.03.030

6. Akaltun İ, Yapça Ö, Ayaydın H, Kara T. An evaluation of attention deficit hyperactivity disorder and specific learning disorder in children born to diabetic gravidas: A case control study. Alpha Psychiatry. 2019;(0):1. doi:10.5455/apd.10445

7. Rowland J, Wilson CA. The association between gestational diabetes and ASD and ADHD: A systematic review and meta-analysis. Sci Rep. 2021;11(1):5136. doi:10.1038/s41598-021-84573-3

8. Nogueira Avelar eSilva R, Yu Y, Liew Z, Vested A, Sørensen HT, Li J. Associations of maternal diabetes during pregnancy with psychiatric disorders in offspring during the first 4 decades of life in a population-based danish birth cohort. JAMA Netw Open. 2021;4(10):e2128005. doi:10.1001/jamanetworkopen.2021.28005

9. Rodolaki K, Pergialiotis V, Iakovidou N, Boutsikou T, Iliodromiti Z, Kanaka-Gantenbein C. The impact of maternal diabetes on the future health and neurodevelopment of the offspring: A review of the evidence. Front Endocrinol. 2023;14:1125628. doi:10.3389/fendo.2023.1125628

10. Kinnunen J, Vääräsmäki M, Keikkala E, et al. Exploring the association between gestational diabetes exposure and mental and behavioural disorders in offspring: The finnish gestational diabetes (FinnGeDi) register-based study. Eur Child Adolesc Psychiatry. 2025;34(12):3987–3998. doi:10.1007/s00787-025-02800-y

11. Ahmed S,, Cano MÁSánchez M, Hu N, Ibañez G. Effect of exposure to maternal diabetes during pregnancy on offspring’s brain cortical thickness and neurocognitive functioning. Child Neuropsychol. 2023;29(4):588–606. doi:10.1080/09297049.2022.2103105

12. Lynch KM, Alves JM, Chow T, et al. Selective morphological and volumetric alterations in the hippocampus of children exposed in utero to gestational diabetes mellitus. Hum Brain Mapp. 2021;42(8):2583–2592. doi:10.1002/hbm.25390

13. Nivins S, Klingberg T. Effects of prenatal exposure to maternal diabetes mellitus on deep grey matter structures and attention deficit hyperactivity disorder symptoms in children. Acta Paediatr. 2023;112(7):1511–1523. doi:10.1111/apa.16756

14. Luo S, Hsu E, Lawrence KE, et al. Associations among prenatal exposure to gestational diabetes mellitus, brain structure, and child adiposity markers. Obesity. 2023;31(11):2699–2708. doi:10.1002/oby.23901

15. Hsu E, Pickering TA, Luo S. Prenatal exposure to gestational diabetes mellitus is associated with greater pre-pubertal BMI growth and faster post-pubertal cortical thinning during peri-adolescence. Pediatr Obes. 2026;21(1):e70069. doi:10.1111/ijpo.70069

16. Page KA, Luo S, Wang X, et al. Children Exposed to Maternal Obesity or Gestational Diabetes Mellitus During Early Fetal Development Have Hypothalamic Alterations That Predict Future Weight Gain. Diabetes Care. 2019;42(8):1473–1480. doi:10.2337/dc18-2581

17. Luo S, Angelo BC, Chow T, et al. Associations Between Exposure to Gestational Diabetes Mellitus In Utero and Daily Energy Intake, Brain Responses to Food Cues, and Adiposity in Children. Diabetes Care. 2021;44(5):1185–1193. doi:10.2337/dc20-3006

18. Zhao S, Semeia L, Veit R, et al. Exposure to gestational diabetes mellitus in utero impacts hippocampal functional connectivity in response to food cues in children. Int J Obes. 2024;48(12):1728–1734. doi:10.1038/s41366-024-01608-1

19. Sappler M, Volleritsch N, Hammerl M, et al. Microstructural Brain Development and Neurodevelopmental Outcome of Very Preterm Infants of Mothers with Gestational Diabetes Mellitus. Neonatology. 2023;120(6):768–775. doi:10.1159/000533335

20. Oikonomou E, Chatzakis C, Stavros S, et al. A review of the impact of gestational diabetes on fetal brain development: An update on neurosonographic markers during the last decade. Life Basel. 2025;15(2):210. doi:10.3390/life15020210

21. Spear LP. The adolescent brain and age-related behavioral manifestations. Neurosci Biobehav Rev. 2000;24(4):417–463. doi:10.1016/s0149-7634(00)00014-2

22. Gogtay N, Giedd JN, Lusk L, et al. Dynamic mapping of human cortical development during childhood through early adulthood. Proc Natl Acad Sci U S A. 2004;101(21):8174–8179. doi:10.1073/pnas.0402680101

23. Shaw P, Kabani NJ, Lerch JP, et al. Neurodevelopmental trajectories of the human cerebral cortex. J Neurosci. 2008;28(14):3586–3594. doi:10.1523/JNEUROSCI.5309-07.2008

24. Young AL, Marinescu RV, Oxtoby NP, et al. Uncovering the heterogeneity and temporal complexity of neurodegenerative diseases with subtype and stage inference. Nat Commun. 2018;9(1):4273. doi:10.1038/s41467-018-05892-0

25. Garavan H, Bartsch H, Conway K, et al. Recruiting the ABCD sample: Design considerations and procedures. Dev Cogn Neurosci. 2018;32:16–22. doi:10.1016/j.dcn.2018.04.004

26. Dick AS, Lopez DA, Watts AL, et al. Meaningful associations in the adolescent brain cognitive development study. NeuroImage. 2021;239:118262. doi:10.1016/j.neuroimage.2021.118262

27. Kahn HS, Divers J, Fino NF, et al. Alternative waist-to-height ratios associated with risk biomarkers in youth with diabetes: Comparative models in the SEARCH for diabetes in youth study. Int J Obes 2005. 2019;43(10):1940–1950. doi:10.1038/s41366-019-0354-8

28. Flegal KM, Cole TJ. Construction of LMS parameters for the centers for disease control and prevention 2000 growth charts. Natl Health Stat Rep. 2013;(63):1–3.

29. Freedman DS, Berenson GS. Tracking of BMI z scores for severe obesity. Pediatrics. 2017;140(3):e20171072. doi:10.1542/peds.2017-1072

30. Sharma AK, Metzger DL, Daymont C, Hadjiyannakis S, Rodd CJ. LMS tables for waist-circumference and waist-height ratio Z-scores in children aged 5–19 y in NHANES III: association with cardio-metabolic risks. Pediatr Res. 2015;78(6):723–729. doi:10.1038/pr.2015.160

31. Denboer JW, Nicholls C, Corte C, Chestnut K. National Institutes of Health Toolbox Cognition Battery. Arch Clin Neuropsychol. 2014;29(7):692–694. doi:10.1093/arclin/acu033

32. Casaletto KB, Umlauf A, Beaumont J, et al. Demographically Corrected Normative Standards for the English Version of the NIH Toolbox Cognition Battery. J Int Neuropsychol Soc JINS. 2015;21(5):378–391. doi:10.1017/S1355617715000351

33. Kaufman J, Birmaher B, Brent D, et al. Schedule for affective disorders and schizophrenia for school-age children-present and lifetime version (K-SADS-PL): Initial reliability and validity data. J Am Acad Child Adolesc Psychiatry. 1997;36(7):980–988. doi:10.1097/00004583-199707000-00021

34. Achenbach TM, Ruffle TM. The child behavior checklist and related forms for assessing behavioral/emotional problems and competencies. Pediatr Rev. 2000;21(8):265–271. doi:10.1542/pir.21-8-265

35. Hagler DJ, Hatton S, Cornejo MD, et al. Image processing and analysis methods for the adolescent brain cognitive development study. NeuroImage. 2019;202:116091. doi:10.1016/j.neuroimage.2019.116091

36. Fortin JP, Parker D, Tunç B, et al. Harmonization of multi-site diffusion tensor imaging data. NeuroImage. 2017;161:149–170. doi:10.1016/j.neuroimage.2017.08.047

37. Fortin JP, Cullen N, Sheline YI, et al. Harmonization of cortical thickness measurements across scanners and sites. NeuroImage. 2018;167:104–120. doi:10.1016/j.neuroimage.2017.11.024

38. Beer JC, Tustison NJ, Cook PA, et al. Longitudinal ComBat: A method for harmonizing longitudinal multi-scanner imaging data. NeuroImage. 2020;220:117129. doi:10.1016/j.neuroimage.2020.117129

39. Kendall MG. A new measure of rank correlation. Biometrika. 1938;30(1-2):81–93. doi:10.1093/biomet/30.1-2.81

40. Bethlehem R a. I, Seidlitz J, White SR, et al. Brain charts for the human lifespan. Nature. 2022;604(7906):525–533. doi:10.1038/s41586-022-04554-y

41. Lawson RG, Jurs PC. New index for clustering tendency and its application to chemical problems. J Chem Inf Comput Sci. 1990;30(1):36–41. doi:10.1021/ci00065a010

42. Gelman A, Hwang J, Vehtari A. Understanding predictive information criteria for bayesian models. Stat Comput. 2013;24(6):997–1016. doi:10.1007/s11222-013-9416-2

43. Bhattacharyya A. On a measure of divergence between two multinomial populations. Sankhyā Indian J Stat 1933-1960. 1946;7(4):401–406.

44. Craig AD. How do you feel? Interoception: The sense of the physiological condition of the body. Nat Rev Neurosci. Published online 2002. doi:10.1038/nrn894

45. Lee SH, Zabolotny JM, Huang H, Lee H, Kim YB. Insulin in the nervous system and the mind: Functions in metabolism, memory, and mood. Mol Metab. 2016;5(8):589–601. doi:10.1016/j.molmet.2016.06.011

46. Simmons WK, DeVille DC. Interoceptive contributions to healthy eating and obesity. Curr Opin Psychol. 2017;17:106–112. doi:10.1016/j.copsyc.2017.07.001

47. Botvinick MM, Braver TS, Barch DM, Carter CS, Cohen JD. Conflict monitoring and cognitive control.Psychol Rev. 2001;108(3):624–652. doi:10.1037/0033-295x.108.3.624

48. Shenhav A, Botvinick MM, Cohen JD. The expected value of control: An integrative theory of anterior cingulate cortex function. Neuron. 2013;79(2):217–240. doi:10.1016/j.neuron.2013.07.007

49. Rolls ET. Taste, olfactory, and food reward value processing in the brain. Prog Neurobiol. 2015;127-128:64–90. doi:10.1016/j.pneurobio.2015.03.002

50. Kevin O, James G. The cognitive control of emotion. Trends Cogn Sci. Published online May 2005. doi:10.1016/j.tics.2005.03.010

51. Arnsten AFT, Li BM. Neurobiology of executive functions: Catecholamine influences on prefrontal cortical functions. Biol Psychiatry. 2005;57(11):1377–1384. doi:10.1016/j.biopsych.2004.08.019

52. Hibar DP, Westlye LT, Doan NT, et al. Cortical abnormalities in bipolar disorder: An MRI analysis of 6503 individuals from the ENIGMA bipolar disorder working group. Mol Psychiatry. 2018;23(4):932–942. doi:10.1038/mp.2017.73

53. Rakic P. A small step for the cell, a giant leap for mankind: a hypothesis of neocortical expansion during evolution. Trends Neurosci. 1995;18(9):383–388. doi:10.1016/0166-2236(95)93934-p

54. Panizzon MS, Fennema-Notestine C, Eyler LT, et al. Distinct genetic influences on cortical surface area and cortical thickness. Cereb Cortex N Y NY. 2009;19(11):2728–2735. doi:10.1093/cercor/bhp026

55. Grasby KL, Jahanshad N, Painter JN, et al. The genetic architecture of the human cerebral cortex.Science. 2020;367(6484):eaay6690. doi:10.1126/science.aay6690

56. Tamnes CK, Østby Y, Fjell AM, Westlye LT, Due-Tønnessen P, Walhovd KB. Brain maturation in adolescence and young adulthood: Regional age-related changes in cortical thickness and white matter volume and microstructure. Cereb Cortex. 2010;20(3):534–548. doi:10.1093/cercor/bhp118

57. Cannon TD, Chung Y, He G, et al. Progressive reduction in cortical thickness as psychosis develops: A multisite longitudinal neuroimaging study of youth at elevated clinical risk. Biol Psychiatry. 2015;77(2):147–157. doi:10.1016/j.biopsych.2014.05.023

58. Ornoy A. Embryonic oxidative stress as a mechanism of teratogenesis with special emphasis on diabetic embryopathy. Reprod Toxicol. 2007;24(1):31–41. doi:10.1016/j.reprotox.2007.04.004

