## Supplemental Table 1 to 7 for "Identification of Heterogeneous Cortical Thickness Patterns Associated with Prenatal Gestational Diabetes Exposure: A SuStaIn-Based Subtyping Study"

*Supplementary Table 1 Demographic, anthropometric, and psychiatric characteristics by GDM status.*

| Variable | GDM (N=573) | Non-GDM (N=2854) | Total (N=3427) | Effect Size | p value |
|---|---|---|---|---|---|
| **Age, months** | 119.02 (7.34) | 118.97 (7.50) | 118.98 (7.47) | -0.01 | 0.899 |
| **Sex (Female)** | 272 (47.5%) | 1458 (51.1%) | 1730 (50.5%) | 0.02 | 0.114 |
| **Race/ethnicity** | | | | 0.1 | < 0.001 |
| White | 342 (59.7%) | 2070 (72.5%) | 2412 (70.4%) | | |
| Black | 78 (13.6%) | 267 (9.4%) | 345 (10.1%) | | |
| Others | 153 (26.7%) | 517 (18.1%) | 670 (19.6%) | | |
| **Family income** | | | | 0.13 | < 0.001 |
| [<50K] | 187 (32.6%) | 580 (20.3%) | 767 (22.4%) | | |
| [>=100K] | 210 (36.6%) | 1503 (52.7%) | 1713 (50.0%) | | |
| [>=50K & <100K] | 176 (30.7%) | 771 (27.0%) | 947 (27.6%) | | |
| **Parental education (bachelor's degree or above)** | 316 (55.1%) | 2027 (71.0%) | 2343 (68.4%) | 0.13 | < 0.001 |
| **Pubertal stage** | | | | 0.1 | < 0.001 |
| Prepuberty | 270 (47.1%) | 1684 (59.0%) | 1954 (57.0%) | | |
| Early puberty | 138 (24.1%) | 629 (22.0%) | 767 (22.4%) | | |
| Mid–post puberty | 165 (28.8%) | 541 (19.0%) | 706 (20.6%) | | |
| **Weight status** | | | | 0.63 | < 0.001 |
| Underweight | 16 (2.8%) | 0 (0.0%) | 16 (0.5%) | | |
| Healthy | 319 (55.7%) | 2854 (100.0%) | 3173 (92.6%) | | |
| Overweight | 103 (18.0%) | 0 (0.0%) | 103 (3.0%) | | |
| Obese | 135 (23.6%) | 0 (0.0%) | 135 (3.9%) | | |
| **BMI** | 19.71 (4.71) | 16.71 (1.44) | 17.21 (2.59) | -1.29 | < 0.001 |
| **BMI-z** | 0.65 (1.22) | -0.09 (0.67) | 0.03 (0.84) | -0.93 | < 0.001 |
| **WHtR** | 0.49 (0.07) | 0.45 (0.04) | 0.45 (0.05) | -1 | < 0.001 |
| **Height(cm)** | 140.99 (7.95) | 139.26 (7.17) | 139.55 (7.33) | -0.24 | < 0.001 |
| **Weight(kg)** | 39.64 (11.93) | 32.57 (4.96) | 33.75 (7.15) | -1.06 | < 0.001 |
| **ADHD**[a] | 36 (6.4%) | 0 (0.0%) | 36 (1.1%) | 0.23 | < 0.001 |
| **Disruptive Behavior Disorder**[a] | 25 (4.4%) | 0 (0.0%) | 25 (0.7%) | 0.19 | < 0.001 |
| **Anxiety**[a] | 66 (11.6%) | 0 (0.0%) | 66 (2.0%) | 0.31 | < 0.001 |
| **OCD**[a] | 42 (7.4%) | 0 (0.0%) | 42 (1.3%) | 0.25 | < 0.001 |
| **MDD/Persistent Depressive Disorder**[a] | 0 (0.0%) | 0 (0.0%) | 0 (0.0%) | | |
| **BD**[a] | 5 (0.9%) | 0 (0.0%) | 5 (0.1%) | 0.08 | < 0.001 |
| **Feeding/Eating Disorder**[a] | 6 (1.1%) | 0 (0.0%) | 6 (0.2%) | 0.09 | < 0.001 |
| **PTSD**[a] | 1 (0.2%) | 0 (0.0%) | 1 (0.0%) | 0.03 | 0.027 |

**Notes**: Values are presented as *n (%)* for categorical variables and *mean (SD)* for continuous variables. Effect sizes are reported as Cohen's d for continuous variables (reference on Subtype 1) and Cramér's V for categorical variables. Effect sizes are rounded to two decimal places and values displayed as 0 indicate very small (non-zero) effects. P values were obtained using two-sample t-tests for continuous variables and chi-square or Fisher's exact tests for categorical variables, as appropriate. P values ≤ 0.05 are shown in bold; + indicates significance after FDR correction. FDR correction was applied separately within demographic, anthropometric, and psychiatric variable groups.

BMI = body mass index; BMI-z = BMI z-score; WHtR = waist-to-height ratio. ADHD = attention-deficit/hyperactivity disorder; OCD = obsessive-compulsive disorder; MDD = major depressive disorder; BD = Bipolar Disorder; ASD = autism spectrum disorder

a.Sample size for these analyses: n = 3,334 (GDM: n = 567; non-GDM: n = 2,767).

*Supplementary Table 2 Psychiatric characteristics with past diagnosis by GDM status.*

| Variable | Subtype 1 (N=322) | Subtype 2 (N=245) | Total (N=567) | Effect Size | p value | pFDR |
|---|---|---|---|---|---|---|
| **ADHD** | 21 (5.8%) | 13 (6.3%) | 34 (6.0%) | 0 | 0.824 | 0.942 |
| **Disruptive Behavior Disorder** | 29 (8.1%) | 17 (8.2%) | 46 (8.1%) | 0 | 0.968 | 0.968 |
| **Anxiety** | 119 (33.1%) | 62 (29.8%) | 181 (31.9%) | 0 | 0.411 | 0.822 |
| **OCD** | 15 (4.2%) | 14 (6.7%) | 29 (5.1%) | 0.04 | 0.184 | 0.735 |
| **MDD/Persistent Depressive Disorder** | 12 (3.4%) | 6 (2.9%) | 18 (3.2%) | 0 | 0.76 | 0.942 |
| **BD** | 19 (5.3%) | 23 (11.1%) | 42 (7.4%) | 0.1 | **0.012** | 0.092 |
| **Feeding/Eating Disorder** | 2 (0.6%) | 0 (0.0%) | 2 (0.4%) | 0.02 | 0.281 | 0.749 |
| **PTSD** | 10 (2.8%) | 5 (2.4%) | 15 (2.6%) | 0 | 0.785 | 0.942 |

**Notes**: Values are presented as *n (%)* for categorical variables and *mean (SD)* for continuous variables. Effect sizes are reported as Cramér's V for categorical variables. Effect sizes are rounded to two decimal places and values displayed as 0 indicate very small (non-zero) effects. ADHD = attention-deficit/hyperactivity disorder; OCD = obsessive-compulsive disorder; MDD = major depressive disorder; BD = bipolar disorder; PTSD = posttraumatic stress disorder.

*Supplementary Table 3 Baseline CBCL comparisons between subtypes.*

| | Subtype 1 (N=362) | Subtype 2 (N=211) | Total (N=573) | Effect Size | p value | pFDR |
|---|---|---|---|---|---|---|
| **Anxious/Depressed** | 53.63 (6.02) | 53.49 (5.81) | 53.58 (5.94) | -0.02 | 0.783 | 0.877 |
| **Withdrawn/Depressed** | 53.20 (5.52) | 53.05 (5.16) | 53.15 (5.39) | -0.03 | 0.737 | 0.877 |
| **Somatic Complaints** | 54.66 (6.13) | 55.43 (6.43) | 54.95 (6.24) | 0.12 | 0.156 | 0.618 |
| **Social Problems** | 52.98 (5.04) | 53.09 (4.81) | 53.02 (4.95) | 0.02 | 0.789 | 0.877 |
| **Thought Problems** | 53.51 (5.46) | 53.99 (5.68) | 53.69 (5.54) | 0.09 | 0.315 | 0.618 |
| **Attention Problems** | 53.29 (5.52) | 54.14 (6.67) | 53.60 (5.98) | 0.14 | 0.102 | 0.618 |
| **Rule-Breaking Behavior** | 52.58 (4.77) | 52.98 (4.98) | 52.73 (4.85) | 0.08 | 0.34 | 0.618 |
| **Aggressive Behavior** | 52.33 (4.91) | 53.02 (5.67) | 52.58 (5.21) | 0.13 | 0.124 | 0.618 |
| **DSM Depression** | 53.27 (5.49) | 53.34 (5.16) | 53.29 (5.37) | 0.01 | 0.878 | 0.898 |
| **DSM Anxiety Disorder** | 53.70 (6.42) | 53.89 (6.52) | 53.77 (6.45) | 0.03 | 0.741 | 0.877 |
| **DSM Somatic Problems** | 55.23 (6.60) | 55.85 (7.04) | 55.46 (6.77) | 0.09 | 0.291 | 0.618 |
| **DSM ADHD** | 52.64 (4.86) | 53.61 (6.31) | 53.00 (5.45) | 0.18 | **0.04** | 0.533 |
| **DSM Oppositional Defiant** | 53.06 (5.06) | 53.96 (5.96) | 53.39 (5.42) | 0.17 | 0.053 | 0.533 |
| **DSM Conduct Disorder** | 52.84 (5.28) | 52.78 (5.04) | 52.82 (5.19) | -0.01 | 0.898 | 0.898 |
| **Sluggish Cognitive Tempo** | 52.67 (4.94) | 52.82 (5.17) | 52.72 (5.02) | 0.03 | 0.736 | 0.877 |
| **Obsessive-Compulsive Problems** | 53.49 (5.91) | 53.35 (5.68) | 53.44 (5.82) | -0.02 | 0.777 | 0.877 |
| **Stress Problems** | 52.83 (5.53) | 53.40 (5.85) | 53.04 (5.65) | 0.10 | 0.245 | 0.618 |
| **Internalizing Problems** | 48.36 (10.48) | 48.99 (10.12) | 48.59 (10.34) | 0.06 | 0.485 | 0.808 |
| **Externalizing Problems** | 44.97 (9.89) | 45.82 (10.66) | 45.28 (10.18) | 0.08 | 0.34 | 0.618 |
| **Total Problems** | 45.59 (10.60) | 46.67 (11.05) | 45.99 (10.77) | 0.10 | 0.248 | 0.618 |

**Notes:** Values are presented as *n (%)* for categorical variables and *Mean (SD)* for continuous variables. Effect sizes are reported as Cohen's d for continuous variables (reference on Subtype 1). Positive *d* values indicate higher values in Subtype 2, and negative values indicate higher values in Subtype 1. Effect sizes are rounded to two decimal places and values displayed as 0 indicate very small (non-zero) effects. Bolded values are statistically significant.

*Supplementary Table 4 Baseline NIH Toolbox Comparisons Between Subtypes.*

| | Subtype 1 (N=362) | Subtype 2 (N=211) | Total (N=573) | Effect Size | p value | pFDR |
|---|---|---|---|---|---|---|
| **Picture Vocabulary** | | | | 0.03 | 0.731 | 0.813 |
| N-Miss | 17 | 19 | 36 | | | |
| Mean (SD) | 52.32 (11.31) | 51.98 (10.13) | 52.20 (10.90) | | | |
| **Oral Reading Recognition** | | | | -0.17 | 0.579 | 0.54 |
| N-Miss | 18 | 20 | 38 | | | |
| Mean (SD) | 48.90 (11.09) | 49.46 (11.02) | 49.10 (11.06) | | | |
| **Flanker Inhibitory Control** | | | | -0.04 | 0.054 | 0.813 |
| N-Miss | 18 | 19 | 37 | | | |
| Mean (SD) | 45.16 (8.43) | 46.68 (9.23) | 45.70 (8.74) | | | |
| **Pattern Comparison Processing Speed** | | | | -0.11 | 0.652 | 0.813 |
| N-Miss | 19 | 19 | 38 | | | |
| Mean (SD) | 44.51 (14.36) | 45.10 (14.87) | 44.73 (14.53) | | | |
| **Picture Sequence Memory** | | | | -0.05 | 0.22 | 0.813 |
| N-Miss | 18 | 19 | 37 | | | |
| Mean (SD) | 49.08 (11.26) | 50.31 (10.96) | 49.52 (11.16) | | | |
| **Dimensional Change Card Sort** | | | | -0.03 | 0.729 | 0.813 |
| N-Miss | 18 | 19 | 37 | | | |
| Mean (SD) | 47.35 (8.84) | 47.64 (9.93) | 47.46 (9.24) | | | |
| **List Sorting Working Memory** | | | | 0.04 | 0.658 | 0.813 |
| N-Miss | 20 | 21 | 41 | | | |
| Mean (SD) | 49.60 (10.33) | 49.20 (9.05) | 49.45 (9.89) | | | |
| **Crystallized Composite** | | | | -0.01 | 0.873 | 0.873 |
| N-Miss | 23 | 20 | 43 | | | |
| Mean (SD) | 50.66 (11.29) | 50.82 (10.62) | 50.72 (11.04) | | | |
| **Fluid Composite** | | | | -0.1 | 0.25 | 0.813 |
| N-Miss | 25 | 21 | 46 | | | |
| Mean (SD) | 45.04 (11.00) | 46.21 (11.58) | 45.46 (11.22) | | | |
| **Total Cognition Composite** | | | | -0.07 | 0.444 | 0.813 |
| N-Miss | 26 | 21 | 47 | | | |
| Mean (SD) | 47.04 (11.24) | 47.81 (10.69) | 47.32 (11.04) | | | |

**Notes:** Values are presented as *n (%)* for categorical variables and *Mean (SD)* for continuous variables. Effect sizes are reported as Cohen's d for continuous variables (reference on Subtype 1). Positive *d* values indicate higher values in Subtype 2, and negative values indicate higher values in Subtype 1. Effect sizes are rounded to two decimal places and values displayed as 0 indicate very small (non-zero) effects. Bolded values are statistically significant.

*Supplementary Table 5 Regional Differences in Brain Morphometry Between Subtype.*

| Variable | Subtype 1 (N=362) | Subtype 2 (N=211) | Total (N=573) | Effect Size | p value | pFDR |
|---|---|---|---|---|---|---|
| ***Cortical Thickness*** | | | | | | |
| Frontal | 2.78 (0.06) | 2.89 (0.06) | 2.82 (0.08) | 1.74 | **< 0.001** | **< 0.001** |
| Parietal | 2.57 (0.07) | 2.67 (0.07) | 2.61 (0.09) | 1.3 | **< 0.001** | **< 0.001** |
| Occipital | 2.09 (0.09) | 2.16 (0.09) | 2.12 (0.10) | 0.79 | **< 0.001** | **< 0.001** |
| Temporal | 2.93 (0.09) | 3.05 (0.08) | 2.98 (0.10) | 1.44 | **< 0.001** | **< 0.001** |
| Cingulate | 2.61 (0.08) | 2.71 (0.08) | 2.65 (0.09) | 1.35 | **< 0.001** | **< 0.001** |
| Insula | 3.11 (0.10) | 3.26 (0.09) | 3.16 (0.12) | 1.64 | **< 0.001** | **< 0.001** |
| ***Brain Volume*** | | | | | | |
| Frontal | 10763.10 (943.92) | 10972.11 (1074.76) | 10840.06 (998.29) | 0.21 | **0.016** | **0.023** |
| Parietal | 14468.66 (1518.79) | 14826.57 (1674.67) | 14600.46 (1585.97) | 0.23 | **0.009** | **0.018** |
| Occipital | 7154.97 (881.04) | 7283.17 (824.49) | 7202.18 (862.14) | 0.15 | 0.086 | 0.103 |
| Temporal | 8413.03 (824.46) | 8684.71 (917.13) | 8513.08 (868.91) | 0.32 | **< 0.001** | **0.002** |
| Cingulate | 3048.85 (365.34) | 3089.24 (389.65) | 3063.72 (374.64) | 0.11 | 0.213 | 0.213 |
| Insula | 7525.38 (796.35) | 7719.95 (850.43) | 7597.03 (821.33) | 0.24 | **0.006** | **0.018** |
| ***Surface Area*** | | | | | | |
| Frontal | 3325.81 (311.48) | 3232.96 (338.22) | 3291.62 (324.40) | -0.29 | **< 0.001** | **0.003** |
| Parietal | 4955.77 (525.70) | 4843.49 (557.86) | 4914.42 (540.01) | -0.21 | **0.016** | **0.019** |
| Occipital | 3051.33 (332.50) | 2967.28 (301.49) | 3020.38 (323.71) | -0.26 | **0.003** | **0.004** |
| Temporal | 2378.53 (236.91) | 2332.43 (256.73) | 2361.55 (245.18) | -0.19 | **0.03** | **0.03** |
| Cingulate | 1002.60 (118.87) | 969.75 (121.86) | 990.50 (120.92) | -0.27 | **0.002** | **0.003** |
| Insula | 2374.39 (249.95) | 2295.09 (255.29) | 2345.19 (254.60) | -0.31 | **< 0.001** | **0.002** |

**Note.** Values are presented as mean (standard deviation). Effect sizes are reported as Cohen's *d* (reference on Subtype 1). Positive *d* values indicate higher values in Subtype 2, and negative values indicate higher values in Subtype 1. Bolded values are statistically significant. Cortical thickness is measured in millimeters (mm), brain volume in cubic millimeters ($mm^3$), and surface area in square millimeters ($mm^2$).

*Supplementary Table 6 Longitudinal associations of subtype and time with CBCL(**observations = 1352**).*

| Outcome | Terms | Estimate | 95% CI | p value | pFDR |
|---|---|---|---|---|---|
| **Anxious/Depressed** | subtype2 | -0.15 | (-1.09, 0.80) | 0.764 | 0.919 |
| | time | -0.28 | (-0.54, -0.01) | **0.044** | 0.105 |
| | interaction | 0.24 | (-0.07, 0.56) | 0.130 | 0.789 |
| **Withdrawn/Depressed** | subtype2 | -0.22 | (-1.13, 0.69) | 0.633 | 0.919 |
| | time | 0 | (-0.26, 0.25) | 0.971 | 0.971 |
| | interaction | 0.31 | (0.01, 0.61) | **0.045** | 0.789 |
| **Somatic Complaints** | subtype2 | 0.65 | (-0.33, 1.64) | 0.192 | 0.737 |
| | time | -0.31 | (-0.60, -0.03) | **0.033** | 0.094 |
| | interaction | 0.12 | (-0.22, 0.46) | 0.484 | 0.862 |
| **Social Problems** | subtype2 | 0.05 | (-0.73, 0.84) | 0.891 | 0.919 |
| | time | -0.22 | (-0.43, -0.00) | 0.050 | 0.105 |
| | interaction | 0.11 | (-0.14, 0.37) | 0.387 | 0.862 |
| **Thought Problems** | subtype2 | 0.29 | (-0.63, 1.22) | 0.536 | 0.919 |
| | time | -0.22 | (-0.48, 0.04) | 0.095 | 0.147 |
| | interaction | 0.05 | (-0.25, 0.35) | 0.758 | 0.862 |
| **Attention Problems** | subtype2 | 0.66 | (-0.28, 1.61) | 0.171 | 0.737 |
| | time | -0.07 | (-0.31, 0.17) | 0.552 | 0.612 |
| | interaction | -0.04 | (-0.32, 0.24) | 0.775 | 0.862 |
| **Rule-Breaking Behavior** | subtype2 | 0.38 | (-0.33, 1.10) | 0.294 | 0.737 |
| | time | -0.24 | (-0.43, -0.05) | **0.014** | 0.074 |
| | interaction | -0.1 | (-0.31, 0.12) | 0.394 | 0.862 |
| **Aggressive Behavior** | subtype2 | 0.63 | (-0.18, 1.44) | 0.130 | 0.737 |
| | time | -0.12 | (-0.33, 0.09) | 0.268 | 0.372 |
| | interaction | 0.01 | (-0.24, 0.25) | 0.957 | 0.957 |
| **DSM Depression** | subtype2 | -0.05 | (-0.97, 0.88) | 0.919 | 0.919 |
| | time | 0.08 | (-0.19, 0.35) | 0.582 | 0.612 |
| | interaction | 0.26 | (-0.06, 0.58) | 0.112 | 0.789 |
| **DSM Anxiety Disorder** | subtype2 | 0.12 | (-0.87, 1.12) | 0.806 | 0.919 |
| | time | -0.24 | (-0.53, 0.04) | 0.087 | 0.147 |
| | interaction | 0.1 | (-0.23, 0.43) | 0.544 | 0.862 |
| **DSM Somatic Problems** | subtype2 | 0.54 | (-0.52, 1.60) | 0.318 | 0.737 |
| | time | -0.38 | (-0.70, -0.07) | 0.018 | 0.074 |
| | interaction | 0.12 | (-0.25, 0.49) | 0.533 | 0.862 |
| **DSM ADHD** | subtype2 | 0.84 | (-0.05, 1.72) | 0.065 | 0.679 |
| | time | 0.12 | (-0.10, 0.34) | 0.279 | 0.372 |
| | interaction | -0.07 | (-0.32, 0.18) | 0.584 | 0.862 |
| **DSM Oppositional Defiant** | subtype2 | 0.77 | (-0.06, 1.60) | 0.068 | 0.679 |
| | time | -0.19 | (-0.41, 0.03) | 0.092 | 0.147 |
| | interaction | -0.04 | (-0.30, 0.21) | 0.738 | 0.862 |
| **DSM Conduct Disorder** | subtype2 | 0.05 | (-0.73, 0.84) | 0.895 | 0.919 |
| | time | -0.29 | (-0.49, -0.09) | **0.005** | 0.074 |
| | interaction | 0.05 | (-0.19, 0.28) | 0.705 | 0.862 |
| **Sluggish Cognitive Tempo** | subtype2 | -0.06 | (-0.88, 0.75) | 0.877 | 0.919 |

| | | | | | |
|---|---|---|---|---|---|
| | time | -0.07 | (-0.30, 0.15) | 0.518 | 0.610 |
| | interaction | 0.17 | (-0.09, 0.44) | 0.197 | 0.789 |
| **Obsessive-Compulsive Problems** | subtype2 | -0.17 | (-1.09, 0.75) | 0.719 | 0.919 |
| | time | -0.25 | (-0.51, 0.00) | 0.052 | 0.105 |
| | interaction | 0.14 | (-0.17, 0.44) | 0.379 | 0.862 |
| **Stress Problems** | subtype2 | 0.46 | (-0.44, 1.36) | 0.320 | 0.737 |
| | time | -0.1 | (-0.35, 0.14) | 0.406 | 0.508 |
| | interaction | 0.04 | (-0.24, 0.32) | 0.780 | 0.862 |
| **Internalizing Problems** | subtype2 | 0.45 | (-1.25, 2.15) | 0.605 | 0.919 |
| | time | -0.55 | (-1.00, -0.09) | **0.019** | 0.074 |
| | interaction | 0.36 | (-0.17, 0.88) | 0.185 | 0.789 |
| **Externalizing Problems** | subtype2 | 0.8 | (-0.81, 2.41) | 0.331 | 0.737 |
| | time | -0.45 | (-0.84, -0.06) | **0.023** | 0.078 |
| | interaction | 0.05 | (-0.40, 0.50) | 0.819 | 0.862 |
| **Total Problems** | subtype2 | 0.81 | (-0.98, 2.60) | 0.376 | 0.752 |
| | time | -0.52 | (-0.95, -0.09) | **0.018** | 0.074 |
| | interaction | 0.11 | (-0.38, 0.61) | 0.658 | 0.862 |

**Notes**: Results from linear mixed-effects models assessing subtype, time, and their interaction on CBCL t-scores, adjusting baseline age, sex, race, household income, parental education, and puberty stage. Bolded values are statistically significant; p stands for raw p value and pFDR stands for p values after FDR correction. ADHD = attention-deficit/hyperactivity disorder.

*Supplementary Table 7 Longitudinal associations of subtype and time with NIH toolbox(observations = 1352).*

| Outcome | Terms | Estimate | 95% CI | p value | pFDR |
|---|---|---|---|---|---|
| **Dimensional Change Card Sort (Fluid)** | subtype2 | 0.32 | (-1.33, 1.96) | 0.706 | 0.853 |
| | time | 0.96 | (0.11, 1.82) | 0.060 | 0.099 |
| | interaction | -1.87 | (-4.67, 0.93) | 0.275 | 0.839 |
| **Flanker Inhibitory Control (Fluid)** | subtype2 | 1.33 | (-0.24, 2.90) | 0.097 | 0.531 |
| | time | 0.64 | (0.09, 1.20) | **0.023** | **0.045** |
| | interaction | -0.59 | (-1.28, 0.10) | 0.094 | 0.598 |
| **List Sorting Working Memory (Fluid)** | subtype2 | -0.37 | (-2.13, 1.39) | 0.682 | 0.853 |
| | time | 0.83 | (0.20, 1.47) | **0.010** | **0.031** |
| | interaction | 0.53 | (-0.14, 1.20) | 0.120 | 0.598 |
| **Pattern Comparison Processing Speed (Fluid)** | subtype2 | 0.37 | (-2.09, 2.83) | 0.768 | 0.853 |
| | time | 4.17 | (3.38, 4.95) | **<0.001** | **<0.001** |
| | interaction | -0.35 | (-1.30, 0.60) | 0.471 | 0.839 |
| **Picture Sequence Memory (Fluid)** | subtype2 | 1.65 | (-0.35, 3.64) | 0.106 | 0.531 |
| | time | 1.5 | (0.85, 2.15) | **<0.001** | **<0.001** |
| | interaction | -0.14 | (-0.93, 0.64) | 0.721 | 0.839 |
| **Fluid Composite** | subtype2 | 1.16 | (-0.85, 3.17) | 0.260 | 0.853 |
| | time | 0 | (-0.01, 0.01) | 0.998 | 0.999 |
| | interaction | -7.5 | (-7.51, -7.49) | 0.791 | 0.839 |
| **Picture Vocabulary (Crystallized)** | subtype2 | -0.13 | (-1.91, 1.65) | 0.884 | 0.884 |
| | time | -0.6 | (-1.07, -0.13) | **0.013** | **0.031** |
| | interaction | 0.13 | (-0.41, 0.68) | 0.631 | 0.839 |
| **Oral Reading Recognition (Crystallized)** | subtype2 | 0.36 | (-1.46, 2.18) | 0.698 | 0.853 |
| | time | 0.16 | (-0.30, 0.63) | 0.491 | 0.614 |
| | interaction | 0.08 | (-0.45, 0.62) | 0.766 | 0.839 |
| **Crystallized Composite** | subtype2 | 0.28 | (-1.60, 2.16) | 0.767 | 0.853 |
| | time | -0.45 | (-1.06, 0.15) | 0.144 | 0.206 |
| | interaction | -0.08 | (-0.86, 0.70) | 0.839 | 0.839 |
| **Total Cognition Composite** | subtype2 | 0.74 | (-1.23, 2.71) | 0.462 | 0.853 |
| | time | 0 | (-0.01, 0.01) | 0.999 | 0.999 |
| | interaction | -11.5 | (-11.51, -11.49) | 0.695 | 0.839 |

**Notes**: Results from linear mixed-effects models assessing subtype, time, and their interaction on NIH toolbox measures, adjusting baseline age, sex, race, household income, parental education, and puberty stage. Bolded values are statistically significant; p stands for raw p value and pFDR stands for p values after FDR correction.

### Figures

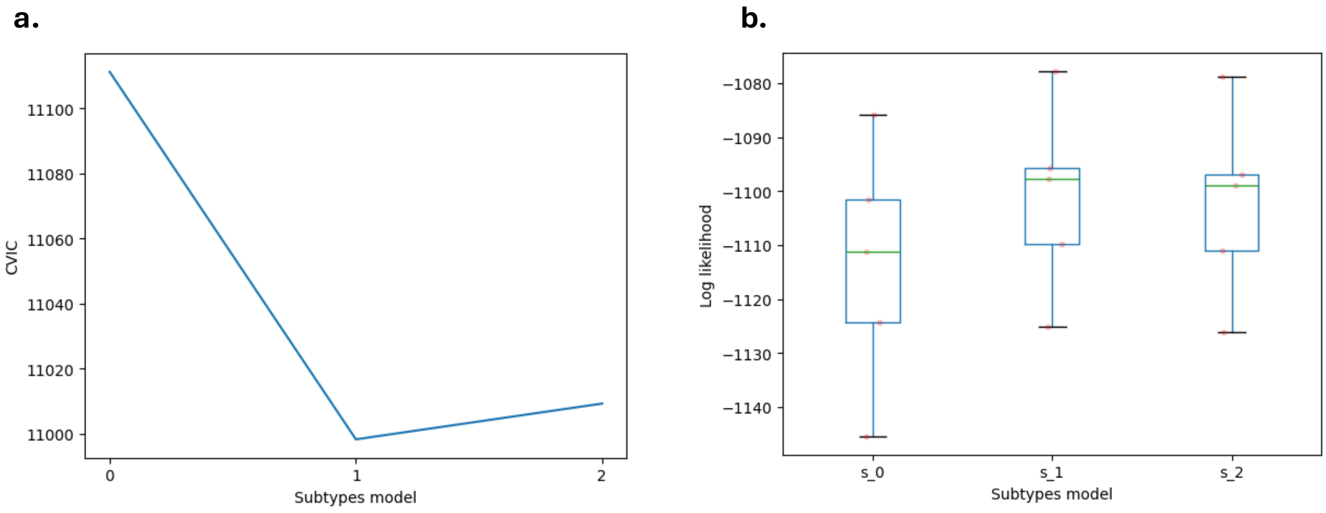


**Supplementary Figure 1. Model selection and cross-validation performance of the SuStaIn model. (a) Cross-validation information criterion (CVIC)** for models with different numbers of subtypes. The lowest CVIC value was observed for the two-subtype model, indicating the optimal balance between model fit and complexity. **(b) Cross-validated log-likelihood distributions** for models with 0, 1, and 2 subtypes ($s_0$, $s_1$, $s_2$). Each boxplot summarizes the log-likelihood values across cross-validation folds, with boxes representing the interquartile range and horizontal lines indicating the median. Individual points correspond to fold-specific estimates. The one- and two-subtype models showed improved likelihood compared with the null model, supporting the presence of heterogeneity in cortical atrophy patterns.

a.

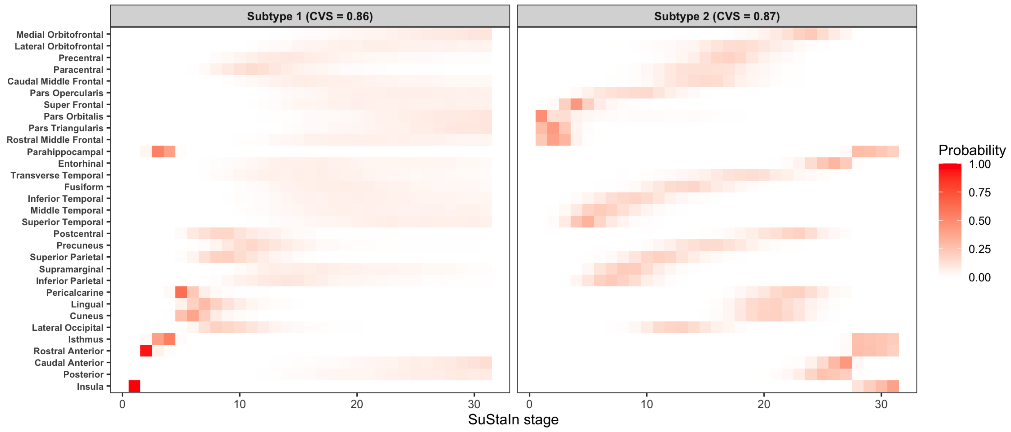


b.

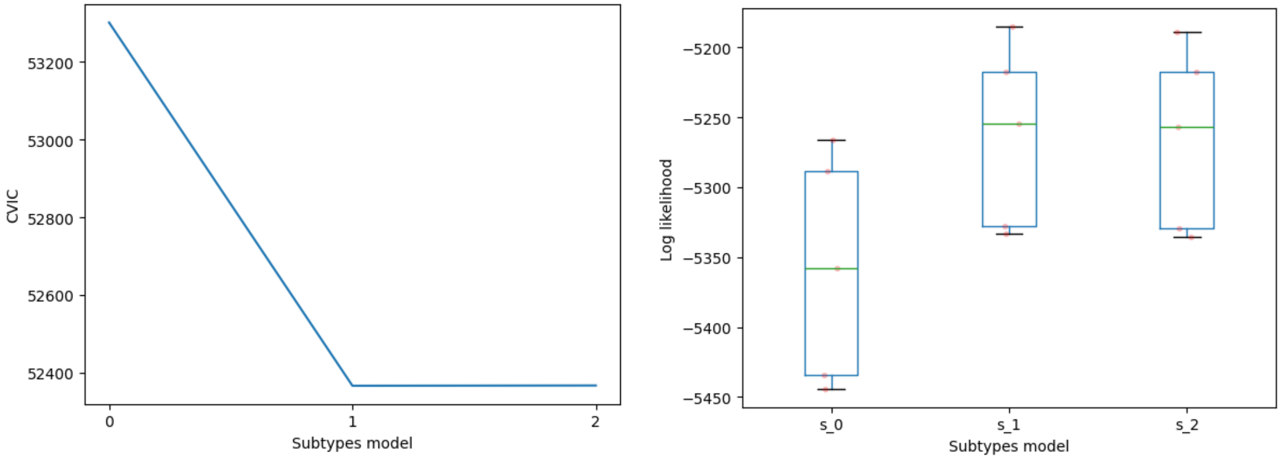


**Supplementary Figure 2. Spatiotemporal patterns cortical thinning identified by the SuStaIn model on ROI-level**. (a) Positional variance diagrams illustrating the inferred progression sequences for the two subtypes identified by the SuStaIn model. Each row represents a cortical ROI, and each column corresponds to a stage in the inferred disease progression. Color intensity indicates the probability that a given region becomes abnormal at a specific stage. Darker colors represent higher probabilities. The CVS values indicate the stability of the subtype sequences across cross-validation folds. (b) Model selection based on the cross-validation information criterion (CVIC). CVIC values are shown for models with different numbers of subtypes. Lower CVIC values indicate better out-of-sample model fit. (c) Distribution of cross-validated log-likelihood values across folds for models with different numbers of subtypes. Boxplots show the variability of log-likelihood across folds, with higher values indicating better model fit.

### Texts

#### Applicability

SuStaIn assumes that structural alterations can be modeled as progressive deviations from a healthy normative mean, enabling the identification of latent heterogeneity in disease trajectories[1–3]. Previous studies have shown that prenatal exposure to gestational diabetes mellitus (GDM) is associated with alterations in offspring brain structure, including reduced global and regional cortical thickness as well as decreased gray matter volume[4,5]. In this study, we selected cortical thickness as the primary biomarker for two main reasons. First, during the developmental window examined in our cohort (baseline age 9–10 years), cortical thickness follows a normative trajectory characterized by gradual decline due to synaptic pruning[6]. This predominantly monotonic decrease aligns well with the linear progression assumption of the SuStaIn framework. In contrast, total cerebral volume and cortical surface area typically peak around early adolescence (approximately 12 years of age), making their developmental trajectories less consistent with this assumption. Second, SuStaIn relies on Markov Chain Monte Carlo (MCMC) estimation, which is computationally intensive. Given our relatively modest sample size of GDM-exposed participants, including a large number of imaging features could reduce model stability. Therefore, we focused on cortical thickness to maintain model robustness while acknowledging that future studies with larger samples could integrate additional structural MRI measures such as gray matter volume.

Building upon previous applications of SuStaIn in neurodevelopmental and psychiatric research[7,8], we conceptualize GDM-related brain alterations not as classical neurodegeneration but rather as an exaggeration of normative neurodevelopmental processes, specifically accelerated cortical thinning. Within this framework, SuStaIn identifies distinct spatiotemporal patterns in which GDM exposure is associated with anatomical deviations that exceed the expected physiological rate of cortical thinning.

1. Fonteijn HM, Modat M, Clarkson MJ, et al. An event-based model for disease progression and its application in familial alzheimer's disease and huntington's disease. *NeuroImage*. 2012;60(3):1880-1889. doi:10.1016/j.neuroimage.2012.01.062

2. Young AL, Oxtoby NP, Daga P, et al. A data-driven model of biomarker changes in sporadic alzheimer's disease. *Brain*. 2014;137(9):2564-2577. doi:10.1093/brain/awu176

3. Young AL, Marinescu RV, Oxtoby NP, et al. Uncovering the heterogeneity and temporal complexity of neurodegenerative diseases with subtype and stage inference. *Nat Commun*. 2018;9(1):4273. doi:10.1038/s41467-018-05892-0

4. Luo S, Hsu E, Lawrence KE, et al. Associations among prenatal exposure to gestational diabetes mellitus, brain structure, and child adiposity markers. *Obesity*. 2023;31(11):2699-2708. doi:10.1002/oby.23901

5. Ahmed S, Cano MÁ, Sánchez M, Hu N, Ibañez G. Effect of exposure to maternal diabetes during pregnancy on offspring's brain cortical thickness and neurocognitive functioning. *Child Neuropsychol*. 2023;29(4):588-606. doi:10.1080/09297049.2022.2103105

6. Bethlehem R a. I, Seidlitz J, White SR, et al. Brain charts for the human lifespan. *Nature*. 2022;604(7906):525-533. doi:10.1038/s41586-022-04554-y

7. Jiang Y, Wang J, Zhou E, et al. Neuroimaging biomarkers define neurophysiological subtypes with distinct trajectories in schizophrenia. *Nat Ment Health*. 2023;1(3):186-199. doi:10.1038/s44220-023-00024-0

8. Jiang Y, Luo C, Wang J, et al. Neurostructural subgroup in 4291 individuals with schizophrenia identified using the subtype and stage inference algorithm. *Nat Commun*. 2024;15:5996. doi:10.1038/s41467-024-50267-3
